# The method of choice to knock-in large inserts via CRISPR

**DOI:** 10.1101/2021.07.01.450700

**Authors:** David Marks, Lisa Bachmann, Lucia Gallego Villarejo, Alexander Geidies, Natalie Heinen, Jannis Anstatt, Thorsten Müller

**Affiliations:** Department of Molecular Biochemistry, Cell Signalling, Ruhr-University Bochum, Bochum, Germany; Institute of Psychiatric Phenomics and Genomics (IPPG), University Hospital, LMU Munich, Munich 80336, Germany

## Abstract

CRISPR/Cas9 gene editing is a revolutionary method used to study gene function by transcript silencing, knock-out, or activation. The knock-in of DNA fragments to endogenous genes of interest is another promising approach to study molecular pathways but is technically challenging. Many approaches have been suggested, but the proof of correct integration has often been relied on less convenient validation experiments. Within this work, we investigated homology-directed repair (HDR), non-homologous end joining (NHEJ), and PCRextension (PCRext) based approaches as three different methods to knock-in large DNA fragments (>1000 bp), and compared feasibility, cost effectiveness, and reliability. As a knock-in fragment, we used a fluorescent reporter sequence in order to directly assess successful integration by microscopy, subsequently proven by sequencing. For NHEJ and PCRext, we demonstrate that it is insufficient to rely on the fluorescent reporter due to false positive results. Both NHEJ and PCRext failed to reliably knock-in large DNA sequences, they were accompanied by massive generation of InDels driving the methodology cost-intensive and non-reliable. In contrast, combination of CRISPR/Cas9 and HDR revealed correct integration, proven by correct fluorescence of the subcellular localization and sequencing, and thus, corresponds to the method of choice for large fragment integration. Next to HEK293T, we demonstrate successful HDR based knock-in in human induced pluripotent stem cells (hiPSCs). Subsequent differentiation of gene-edited hiPSCs into cerebral organoids showed relevance of the approach to study subcellular protein localization and abundance in 3D tissue.

## Introduction

CRISPR/Cas9 gene editing has revolutionized the manipulation of DNA to knock-out genes, activate or silence targets, introduce mutations or protein tags (**Bukhari & Müller, 2019**). What all these methods have in common is their starting point using a short RNA, which recognizes a specific DNA sequence in the genome and later on guides endogenous proteins to the locus of action, where the Cas9 endonuclease induces a double strand-break 3 bp upstream of the protospacer adjacent motif (PAM). According to this function, the short RNA is termed single guide RNA (sgRNA) and the identification of a well-working sequence is the first highly important step in gene editing. A variety of in-silico tools (for example i. CHOPCHOP, https://chopchop.cbu.uib.no/; ii. IDT Design tool https://eu.idtdna.com/site/order/designtool/index/CRISPR_PREDESIGN, iii. CHRISPOR http://crispor.tefor.net/) has been developed to design the required sgRNAs in order to target a specific gene of interest. All these tools aim to optimize the sgRNA quality in terms of low off-target binding sites and rate the sgRNA according to a quality score. Upon in silico identification, the sgRNAs need to be tested in the laboratory towards their cleavage efficiency and specificity. Therefore, the sgRNA is administered to the cell together with the Cas9 enzyme, either as a ribonucleoprotein (RNP) complex or by respective expression vectors. The genomic region containing the sgRNA target site is then PCR amplified and tested. So far, the method of choice is based on a mismatch-specific endonuclease assay, like Surveyor™ nuclease or T71E endonuclease assay (**Vouillot et al., 2015**). In brief, the nuclease cleaves the amplified DNA fragment at the heteroduplex (hybrid of error-prone repaired fragment and the wt fragment) mismatch (**Vouillot et al., 2015**), and afterwards, the fragments are analyzed by gel-electrophoresis. Thereby, the ratio of wildtype (one fragment, no cleavage) and mutation (two shorter fragments, mismatch cleavage) can be determined, thus evaluating the efficiency of the sgRNA. This method is simple in performance, but expensive, the detection limit is low, and its efficiency is limited. Upon identification of proper sgRNAs, the gene editing method to knock-in a large fragment needs to be defined, e.g. in order to monitor proteins by fluorescence tagging (like GFP), to generate new fusion proteins, or to label a cell line (**Arbab et al., 2015; He et al., 2016; Koch et al., 2018; Verma et al., 2017**). So far three different strategies have been reported including the PCR extension (PCRext) method, homology independent targeted integration (HITI) relying on the non-homologous end joining (NHEJ) repair pathway, and homology-directed repair (HDR). PCRext, initially introduced in 2015, has been suggested as a cloning-free and less time-consuming method (**Arbab et al., 2015**). In brief, the desired DNA knock-in fragment is amplified by PCR, at the same time adding short homologous regions (up to 120 bp) by use of extended primers on both ends, which match to the knock-in locus in the genome, up- and downstream of the sgRNA/Cas9 cleavage site. Mammalian cells can then be transfected with the sgRNA, Cas9 and the PCR-generated linear donor template. An alternative strategy is the NHEJ-based knock-in of large DNA fragments. Next to HDR, NHEJ is one of two major repair mechanisms of the cell. NHEJ is (compared to HDR) the more abundant of the two cellular repair mechanisms as it occurs in both proliferating and non-proliferating cells (**Lieber, 2010**), but it is also known to be error-prone (**Suzuki & Izpisua Belmonte, 2018**). In 2016, Suzuki et al. developed a new strategy for the NHEJ-based knock-in of genes that was able to exploit the advantages of NHEJ while overcoming its deficiencies, which they called homology-independent targeted integration (HITI) (**Suzuki et al., 2016**). Upon induction of a double-strand break (DSB) by sgRNA/Cas9, a linearized donor DNA fragment (e.g. as a cleavage product of a plasmid) is integrated via the NHEJ repair pathway ligating the blunt ends of the DSB and the donor DNA. The significant advantage of the HITI strategy is given by the fact that only if the donor DNA is inserted in the correct orientation, it is protected from further cleavage by the Cas9 nuclease. If the donor is reversely integrated or not integrated at all, the sgRNA target sites remain intact leading to re-cleavage. HDR-based methods have been used to achieve knock-ins of DNA fragments for a long time. Similar to NHEJ, the HDR pathway is initiated upon recognition of DSBs. Within the cell, there is a balance between the competing DSB repair mechanisms which is influenced by not completely understood events in the chromatin that include the interaction of multiple proteins, causing HDR to be more prevalent during S/G2 phase, when the sister chromatid is close and available as a recombination partner (**Yang et al., 2020**). The HDR repair machinery is initiated by the nucleolytic degradation of the 5’→3’ strands (at the DSB site) to yield 3’→5’ single stranded DNA (ssDNA) overhangs onto which then the Rad51 recombinase is able to assemble. This structure is invading homologous DNA strands (e.g. the sister chromatid) using them as a template for the repair mechanism (**Chapman et al., 2012**). Thus, providing a DNA repair template (such as an introduced single-stranded oligo DNA nucleotide or dsDNA with the structure: 5’ homology arm - knock-in fragment - 3’ homology arm) allows the knock-in of DNA fragments (**Dickinson et al., 2013**). This technique has been successfully used to induce knock-ins in different mouse models (**Wang et al., 2015**). However, both embryonic stem cells and human induced pluripotent stem cells have been shown to display a particularly low efficiency in HDR-induced knock-in, although the reasons remain unclear (**He et al., 2016**). Here, we provide an in-depth comparison of the three large fragment knock-in strategies PCRext, NHEJ and HDR. Success of knock-in is compared by immunofluorescence analysis of inserted fluorescent proteins as well as sequence analysis. We demonstrate results in HEK293T cells and human induced pluripotent stem cells (hiPSCs), the latter being not affected by the editing approach in respect to their pluripotency.

## Material and Methods

### Cell culture and transfection

HEK293T cells were cultured in Dulbecco’s Modified Eagle Medium (DMEM, Sigma-Aldrich) with 10 % FBS, 1 % Pen/Strep and 1 % L‑glutamine on 100 mm culture dishes. Cells were transfected using the K4® system (Biontex, Germany) according to the manufacturer’s instructions. HiPSCs were cultured in StemFlexTM (Gibco, Thermo Fisher Scientific) on 35 mm dishes (37 °C, 5 % CO_2_) coated with Geltrex^□^ (Gibco, Thermo Fisher Scientific) according to manufacturer’s instructions, and were passaged regularly using ReLeSR (STEMCELL Technologies). For nucleofection, cells were detached with TrypLE™ (Gibco, Thermo Fisher Scientific) before they reached 70 % confluence and counted in order to use 200.000 cells per nucleofection. Equal amounts of tracrRNA and crRNA (Integrated DNA Technologies) were mixed (50 μM) and incubated at 95 °C to form the sgRNA complex. The ribonucleoprotein complex was formed by incubation of the sgRNA complex with the Cas 9 recombinant protein (4:4,8 μM) (Integrated DNA Technologies) at RT. The nucleofector solution and nucleofector supplement (P3 Primary Cell Kit, Lonza) were added to the cells, mixed, and then the electroporation enhancer (Integrated DNA Technologies) was added, followed by the sgRNA complex, Cas9 and 0.8 μg of the HDR-Plasmid (FE65-EGFP; LMNB1-mTagRFP-T, Addgene #114403). The nucleofection was performed using the 4D-Nucleofector™ X Unit (Lonza, Germany) with the CB-150 pulse according to manufacturer’s instructions. Afterwards, cells were cultivated in StemFlex™ cell culture medium supplemented with ROCK inhibitor (STEMCELL Technologies) and HDR enhancer (Integrated DNA Technologies).

### Design of sgRNAs in silico and cloning into MLM vector

The sgRNAs targeting the N- or C-terminus of the gene of interest (FE65, APP, LMNB1) were designed using the web tool CHOPCHOP and were chosen according to the highest score. Multiple sgRNAs were selected. For FE65, all sgRNAs were located in exon 19, in close proximity to the stop codon to enable a C-terminal knock-in of a fluorescent protein coding sequence, and for APP, the sgRNAs are targeting the sequence near the stop codon in exon 18. This strategy was later adapted to obtain the sgRNAs targeting the N-terminal end of the lamin B1 gene. Forward and reverse primers (oligos) consistent for each sgRNA were purchased (Merck, Sigma-Aldrich) including sticky ends corresponding to the cleavage pattern of the restriction enzyme BsmBI according to the protocol of Kwart et al. (**Kwart et al., 2017**).

The oligos were annealed and then cloned into an MLM3636 vector that was digested with BsmBI and linearized (Addgene, #43860). Subsequently, the constructs were transformed into bacteria, followed by plasmid isolation (NEB, Monarch Plasmid Miniprep Kit). The successful cloning was verified via Sanger sequencing. 5’→3’ sgRNA sequences were as follows: FE65 sgRNA 1 GGTATGGGCCCCCAGCCGTT, FE65 sgRNA 2 TGGGGCCCAACACAAGCAGG, FE65 sgRNA 3 GGGGTCTAGAGGCTATTCCT, FE65 sgRNA 4 GTAGGGTGGACTGTCCGCAG, FE65 sgRNA 5 CCCCTGCTGAGTCTGTGGCA, FE65 sgRNA 6 ATGCCCCTCCCCAGTAGCTA, APP sgRNA1 HDR ACAAGTTCTTTGAGCAGATG, APP sgRNA2 HDR ACCTACAAGTTCTTTGAGCA, Lamin sgRNA HDR GGGGTCGCAGTCGCCATGGC. Designed sgRNAs were tested for successful cleavage using the Surveyor Mutation Detection Kit (Integrated DNA Technologies) according to manufacturer’s instructions.

### Knock-in approaches

#### PCR extension approach

To synthesize the donor template, EGFP (Supplementary Figure 1) was amplified and simultaneously flanked with homology arms homologous to the target gene in order to enable the genomic integration. Therefore, long primers (90 bp in total, Sigma-Aldrich) were used, which included an overhang of 30 bp (underlined sequence) and 60 bp of APP homologous region APP scCRISPR 1 fwd AAGATGCAGCAGAACGGCTACGAAAATCCAACCTACAAGTTCTTTGAGCAGATGCAGAA C<u>ATGGAGAGCGACGAGAGCGGCCTGCCCGCC</u>, APP scCRISPR 1rev TACAGCACAGCTGTCAAAAGGCGATAATGAGTAAATCATAAAACGGGTTTGTTTCTTCC C<u>CTAGCGAGATCCGGTGGAGCCGGGTCCGGC</u>. Homology arms were designed to match the sequence up- and downstream of the APP stop codon, while omitting 84 bp at the 3’ site containing the sgRNA binding site (to prevent further double strand breaks through Cas9 after successful HDR) and the stop codon. The PCR fragment was further used to elongate the homology arms to a desired length of 180 bp on either side. Therefore, two additional PCRs with long extension primers were performed, each adding additional 60 bp to the homology arms (APP scCRISPR fw 2 AGCTCTCCTCTTGTTTTTCAGGTTGACGCCGCTGTCACCCCAGAGGAGCGCCACCTGT CC<u>AAGATGCAGCAGAACGGCTACGAAAATCCA</u>, APP scCRISPR 2 rev AGAGAGATAGAATACATTACTGATGTGTGGATTAATTCAAGTTCAGGCATCTACTTGTGT <u>TACAGCACAGCTGTCAAAAGGCGATAATGA</u>). After each elongation, the fragment was analyzed by gel electrophoresis, isolated, purified using a NucleoSpin Gel and PCR Clean‑up Kit (Machery-Nagel, Germany), and sequenced. Finally, the donor template was transfected (K4® Transfection System, Biontex, Germany) in HEK293T cells together with a Cas9 plasmid and the APPext sgRNAs 3 or 21 (APPext sgRNA 3 TTGCTTCACTACCCATCGGT, APPext sgRNA 21 GTCCATTTATAGAATAATGT) to induce the desired EGFP knock-in. Genomic DNA of the transfected cells was isolated using the NucleoSpin DNA RapidLyse (Machery-Nagel, Germany) and the edited regions were PCR-amplified (Supplementary Figure 2). Either the whole region was amplified with primers binding within the homology arms (HA upstream CATTGCACTGGGTTTATCCACTTACTG, HA downstream CAAAGACCCAAAGATACGTGGACAAAA), or one primer was binding within the homology arm, while the second targeted the EGFP gene (HA upstream with EGFP rev TGGTAGAAGCCGTAGCCCATC or HA downstream with EGFP for GGTGGACAGCCACATGC). After gel electrophoresis separation, the band with the expected fragment size was isolated with the NucleoSpin Gel and PCR Clean‑up Kit (Machery-Nagel, Germany) and Sanger sequenced.

#### NHEJ approach

The donor vector design followed the method described by Suzuki et al. (Suzuki et al., 2016) with slight modifications (Figure 2A). Besides the mNectarine tag for integration, a selection cassette comprising an EGFP gene, an IRES site and a Bleomycin resistance (BleoR) controlled by a CMV promoter was included, allowing the selection of positively knocked-in cells via FACS and antibiotic exposure. To be able to further verify the successful knock-in as well as transfection of the cells, the donor vector was provided with a BFP cassette. The approach enables generation of a linear donor fragment within the cell upon sgRNA-mediated cleavage at the two loci *sgRNA donor start* and *sgRNA donor end* (Figure 2A). The BFP signal vanishes after efficient cleavage by Cas9 as it is excluded from the donor fragment that will be inserted.

**Figure 1.**
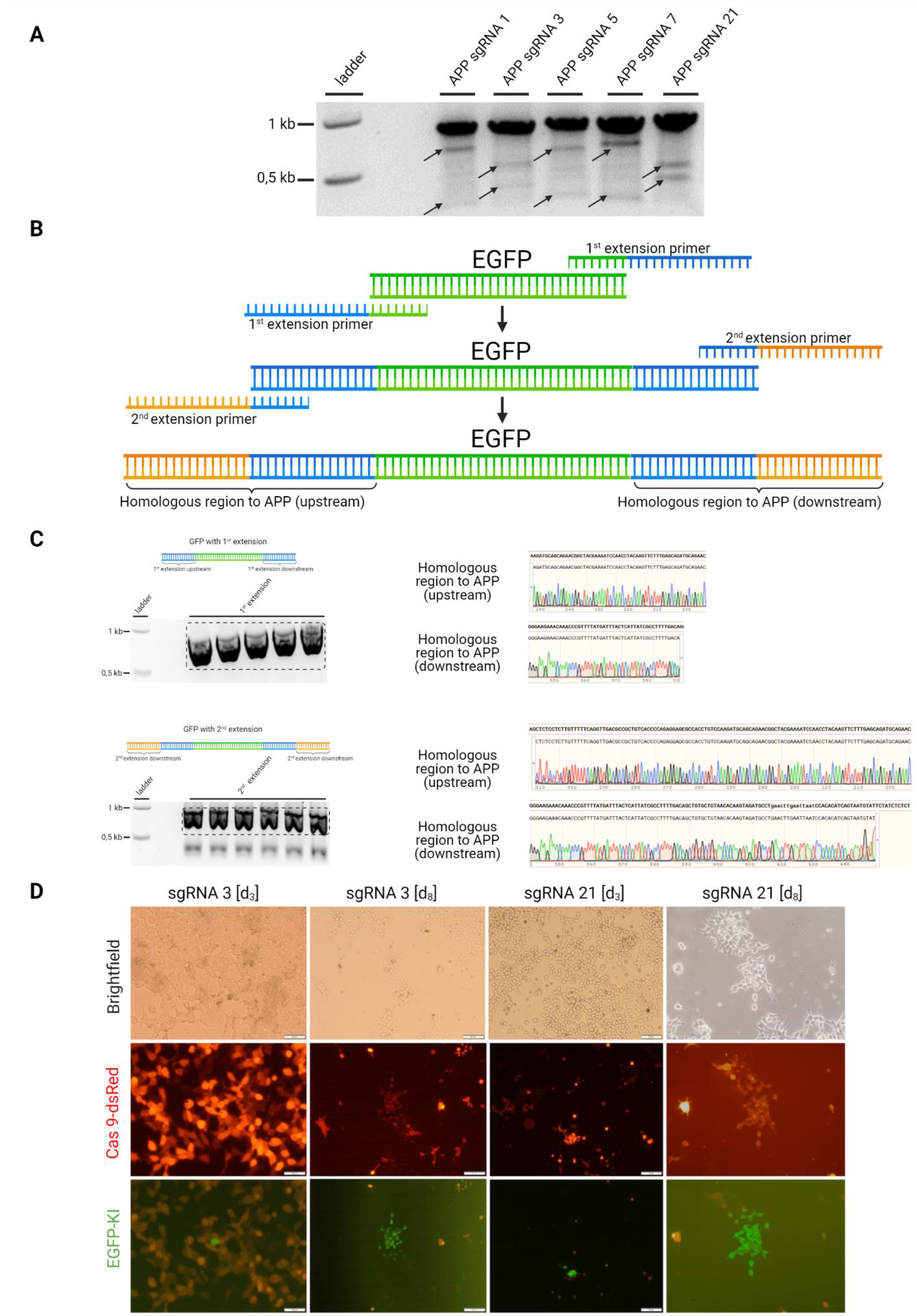
PCRext-driven KI revealed fluorescent cells but failed to show sequencing validation. **(A)** A surveyor™ assay demonstrated high cleavage-efficiency of all sgRNAs tested within this work, which is exemplarily shown for APP. SgRNA numbers (1, 3, 5, 7, 21) represent the CHOPCHOP ranking for off-targets, with a lower number indicating less off-Targets for the respective sgRNA. (B) Two sequential PCR amplifications with extension primers were done, each adding an overhang of 60 bases. The resulting PCR fragment consists of the EGFP with two flanked homologous regions serving as donor template for the CRISPR/Cas9 KI. **(B)** Gel electrophoresis of PCR fragments and sequencing revealed correct extension of the EGFP sequence. **(C)** HEK293T cells transfected with a Cas9 encoding vector, the donor PCRext template, and either sgRNA 3 or sgRNA 21 demonstrated successful transfection (Cas9-dsRed) and suggested correct donor integration (EGFP-KI) at day 3 (d3). Cells were able to proliferate while keeping their green signal (d8). After 14 days, genomic DNA was extracted and multiple PCRs were done in order to amplify the EGFP fragment using various primer binding sites (Supplement Figure 2). While PCR demonstrated products of the expected size for some primer combinations, all sequencing work failed to reveal a correct knock-in suggesting unspecific integration of the EGFP sequence. (Figure created with BioRender.com)

**Figure 2:**
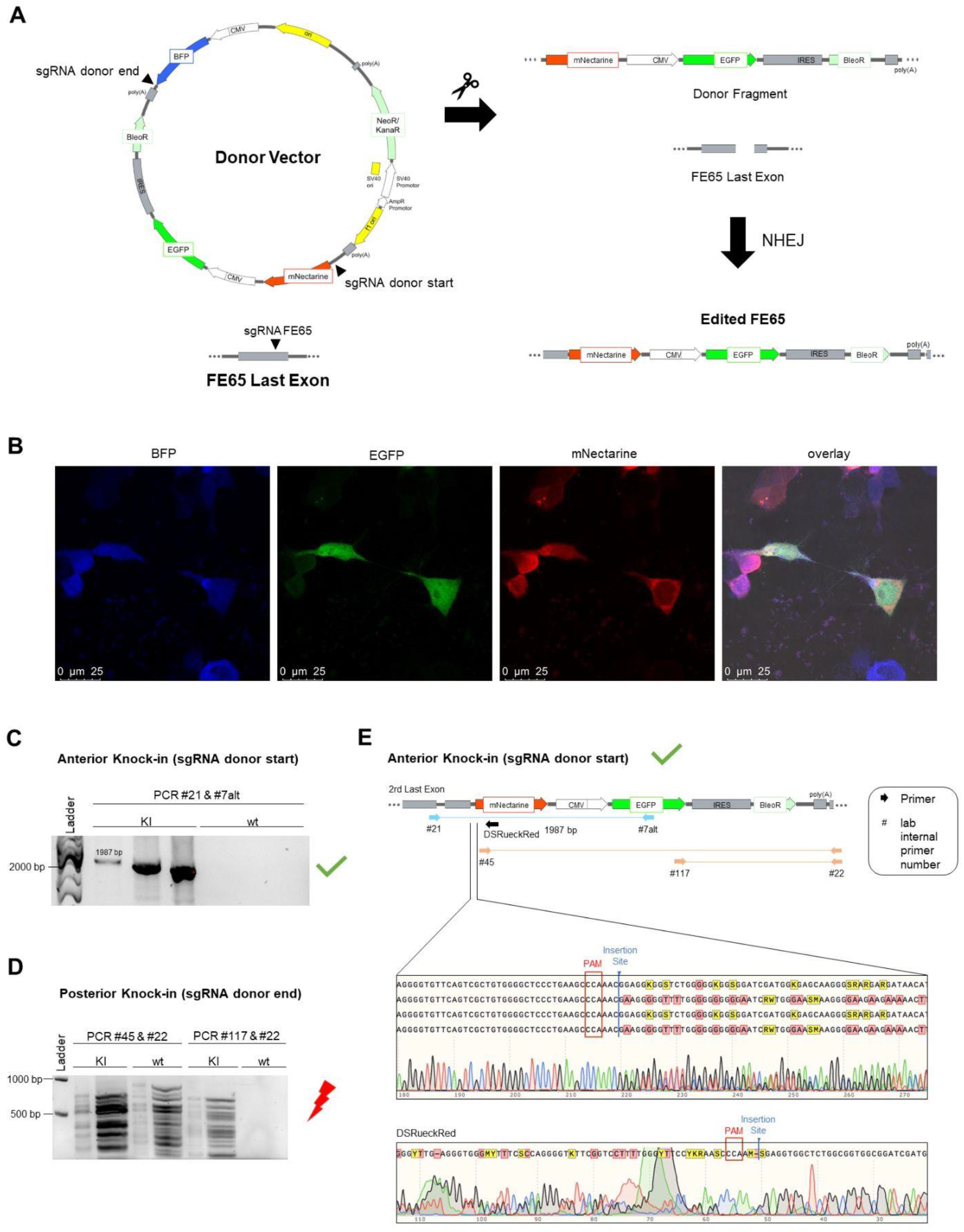
NHEJ-mediated approach demonstrated positive knock-in using immunofluorescence analysis and partial validation by sequencing. **(A)** The donor vector was designed to include an EGFP gene and a Bleomycin resistance (BleoR) regulated by a CMV promoter and an IRES site to allow selection of positively edited cells via FACS and antibiotic exposure. A BFP gene was integrated to verify successful transfection. A first sgRNA (sgRNA donor start) targets the linker region preceding the mNectarine gene, a second sgRNA (sgRNA donor end) leads to a cut behind the polyA motif of the selection cassette. Thus, cleavage by Cas9 creates a fragment comprising the mNectarine gene as well as the selection cassette consisting of EGFP, IRES and BleoR. A third sgRNA (sgRNA FE65) targets the last exon of the gene of interest, consequently creating an insertion site for the excised insertion fragment. NHEJ repair system is integrating the fragment into the target site. **(B)** Transfected HEK293T cells revealed positive fluorescence for BFP, mNectarine, and EGFP suggesting successful integration. Cytosolic localization of the mNectarine signal suggested specific integration. **(C)** Regions surrounding the anterior 5’ (primer #21 & #7alt) and the posterior 3’ (primer #45 & #22 and #117 & #22) knock-in sites were amplified via PCR. Gel electrophoresis analysis surrounding the insertion site anterior (5’) to the knock-in region yielded a clear signal at the expected size of 1978 bp, further verified by the absence of the signal in the wt PCR approach. **(D)** Gel electrophoresis analysis of the PCR fragment surrounding the posterior (3’) insertion site did not yield a distinctive signal indicating an unsuccessful incorporation at the cut site of sgRNA donor end. **(E)** Sequencing using the respective PCR primers demonstrated the expected sequence up to the insertion site, after which the signal becomes indistinct due to the presence of mismatches. However, additional sequencing of the PCR fragment using a reverse primer (#DSRueckRed) binding within the mNectarine fragment confirmed the presence of correctly incorporated mNectarine knock-in fragments (not shown).

#### HDR approach

The backbone vector pUC57-GFP (Addgene #97090) was linearized by PCR and 1000 bp homologous to the FE65 gene (right before and after the STOP codon; including a mutation in the PAM sequence to avoid repeated cleavage after successful knock-in by CRISPR/Cas9) were included. The upstream and downstream homology arms were purchased as gBlocks (Integrated DNA Technologies) including a 15 bp homology sequence to the backbone vector in both sites (3’ and 5’ of the sequence) for In-Fusion® cloning. The In-Fusion® reaction was performed using the In-Fusion® HD Cloning Plus Kit (Takara Bio, Japan) following the manufacturer’s protocol. The plasmid for tagging the N-terminus of the human lamin B1 gene (LMNB1) with RFP was purchased from Addgene (LMNB1-mTagRFP-T, #114403). The plasmid sequence was validated by Sanger sequencing.

#### Knock-in analysis

DNA from transfected HEK293T cells or hiPSCs was isolated to verify the knock-in in the genes of interest. To verify the insertion of the RFP tag N-terminal to LMNB1, a set of primers (LMNB HA Upstream 1 GCCGTGCATTTGGAACCT; LMNB HA downstream II CCTGGTCTACTATCTGCACA) was used to amplify a 2.8 kbp fragment containing a region larger than the two homology arms of 1 kbp length and the RFP. This fragment was then used as template to produce a DNA fragment containing the transition of the RFP protein to the upstream homology arm (primers: LMNB RFP rev GTATGTGGTCTTGAAGTTGC, LMNB_RFP_HA_fw CAAGGGCCAGATTTTAAATTTACAG). The fragment corresponding to the transition of the RFP into the downstream homology arm was prepared using the primers LMNB RFP fw CAACACCGAGATGCTGTA and LMNB HA downstream II CCTGGTCTACTATCTGCACA. For the proof of successful knock-in of EGFP into the FE65 gene, primers binding outside of the homology arms were designed (seq_FE65_1_PreHA1 GAGAGAGCATGATTTGGGGAAATC; seq_FE65_1_PostHA2 GACATGCAACTTTTCCAGAC). The fragment of 2.8 kbp containing a region larger than both the homology arms and the GFP insert was used as a template for further amplification of shorter fragments containing the transitions of the EGFP into the homology arms (GFP-fwd AAGCAGCACGACTTCTTCAAG; sequenc_FE65_2_AS TACTGGGGAGGGGCATAT and GFP-rev CCGTCGTCCTTGAAGAAGATG; qPCR FE65 Forward TAGATGTGATTAATGGGGCCCTCG). The transition of the genome to the homology arms of the knock-in has been validated by sequencing using the primers for the 2.8 kbp fragment amplification.

### Microscopy

Cells were routinely imaged using the Leica DMI 4000B microscope system to assess cell growth. For imaging and observation of transfected and living HEK293T and hiPSCs, an Olympus IX51 inverted microscope system was used, and for confocal imaging, a Leica TCS SP8 confocal microscope system was used. Therefore, cells were grown on coated (0.01 % poly-l-ornithine (Sigma-Aldrich)) glass bottom dishes (ibidi) and imaged with a 63x water objective (HP PL APO CS2 63x/1.2 WATER). Cerebral organoid cryosections were imaged using a Keyence BZ-X fluorescent microscope.

### Cerebral organoids

HiPSCs containing the RFP knock-in for LMNB1 were used to generate cerebral organoids according to the protocol of Lancaster and Knoblich (**Lancaster & Knoblich, 2014**) with minor modifications as described before (**Marks et al., 2021**). Briefly, 9,000 hiPSCs per well were seeded in a round bottom ultra-low attachment 96-well microplate (Corning) with 150 μl of hESC medium supplemented with 4 ng/ml bFGF (Peprotech) and 50 μM ROCK inhibitor (STEMCELL Technologies) in order to form EBs. Medium was exchanged every other day with fresh hESC medium. On day 5 or 6, medium was changed to NI medium and exchanged every other day with fresh NI medium. On day 11 or 12, EBs were embedded in GFR Matrigel (Corning) and placed in a well of a 6-well plate with 3 ml DM-A medium. After incubation for 3-4 days at 37 °C and 5 % CO_2_, the medium was exchanged with 3 ml DM+A medium and the plate was placed on an orbital shaker kept at 37 °C and 5 % CO_2_ for further culture with medium changes every 3-4 days.

## Results

To create the prerequisites for the comparison of the different knock-in approaches, we designed several sgRNAs for the genes of interest FE65, APP, and lamin B1 using the web tool CHOPCHOP. APP and FE65 are of central relevance for the pathophysiology of Alzheimer’s disease in terms of their subcellular localization, abundance and interaction. In addition, we chose lamin B1 for the extension of our approach from HEK293T to hiPSCs, as this protein is of high abundance and reveals a distinct perinuclear localization. Under the assumption that the knock-in is not affecting the protein functionality, the successful knock-in would enable the investigation of FE65- and/or APP-relevant pathways and LMNB1 localization at endogenous expression levels. Surveyor™-assay demonstrated comparable cleavage efficiencies for all studied sgRNAs (Figure 1A, exemplarily shown for APP), as evident by cleaved fragments (Figure 1A, arrows) due to the digestion of the mismatch hybridization.

### Knock-in via PCR extension (PCRext) revealed fluorescent cells, but failed to show sequence validation

As a first attempt, we aimed to apply the PCRext method to tag the APP C-terminal with a coding sequence for EGFP. Extension primers (90 bp in total) were designed, which each added 60 bp homology sequences to each site of the EGFP fragment (Figure 1B). An additional PCR then extended the homology regions at each site to 120 bp. Gel electrophoresis and Sanger sequencing revealed successful elongation of EGFP (Figure 1C). Transfection of HEK293T cells with Cas9-dsRed (to monitor transfection efficiency), the EGFP donor fragment, and either sgRNA 3 or sgRNA 21 revealed successful transfection of Cas9-dsRed (dsRed), while less than 10 % of cells were positive for the knocked-in EGFP. GFP-positive cells were successfully dividing and proliferating into small colonies after several days of incubation (d_8_), suggesting a stable integration of the EGFP gene. Control cells omitting the sgRNA revealed no green fluorescence signal (not shown). To confirm the knock-in at the region of interest, multiple PCRs were performed (Supplementary Figure 2) with primers binding either inside the homologous region to APP (Supplementary Figure 2A), inside the upstream homologous region and EGFP (Supplementary Figure 2B), or inside the downstream homologous region and EGFP (Supplementary Figure 2C). Fragments with the desired sizes could be generated often accompanied by additional signals. However, all subsequent sequencing analyses failed to prove the knock-in and, if successful, the sequence always revealed the wt sequence not supporting an incorporated fragment, pointing to unspecific integration of the EGFP cassette.

### The NHEJ-mediated approach revealed positive knock-in using immunofluorescence analysis and partial validation by sequencing

The designed donor vector (Figure 2A) was transfected together with Cas9 and MLM vectors encoding for *donor start*, *donor end*, and *sgRNA for the target gene,* enabling generation of the linear donor fragment within the cell. Transfected cells revealed positive fluorescence for BFP (transfection control marker), mNectarine (KI marker), and EGFP (KI co-marker). While EGFP was located throughout the cell, mNectarine revealed a cytosolic enrichment (Figure 2B), suggesting successful labeling of the gene of interest. To verify the knock-in, genomic DNA was isolated, the region of interest was PCR-amplified, analyzed via gel electrophoresis and sequenced. The PCR amplification of the 5’ insertion site (anterior knock-in using primers #21 and #7alt) yielded a fragment of the expected size, indicating correct integration upstream of the mNectarine sequence (Figure 2C). In contrast, although various primer combinations were used, amplification of the posterior knock-in sequence was not possible, suggesting a failed knock-in at the 3’ insertion site (Figure 2D). Sequencing analysis for the anterior knock-in site (sequencing primer #21) showed correct alignment with the expected product up to the cut site of *sgRNA donor start* (Figure 2E). Afterwards the chromatogram signal becomes inaccurate including a number of mismatches. Sequencing of the same PCR product using a reverse primer binding within the mNectarine gene (DSRueckRed; primer location in Figure 2E), demonstrated the presence of correctly incorporated fragments, leading to the assumption that the knock-in was successful but includes InDels. Sequencing analysis of the posterior knock-in site failed as expected since no robust PCR fragment of the intended size was amplified.

### HDR as a fast, reliable method to knock-in large fragments

As a final strategy for the KI of a large genomic fragment, we used an HDR-based method relying on a knock-in fragment with long (1000 bp) homology arms flanking the GFP gene. Using InFusion cloning, cassettes for the two homology arms (ordered as gBlocks), EGFP, and the backbone were cloned to establish the donor vector (Figure 3A). 72 h post transfection, PCR amplification (using the same primer pairs as for NHEJ) of isolated DNA revealed fragments of the expected size (~1.4 kbp and ~1.1 kbp) for both the anterior and posterior KI fragments (Figure 3B). Although a (non-sorted) mixture of cells was used, no InDels could be observed, instead sequencing results confirmed correct integration at both seams (Figure 3B). After this validation of successful integration, we applied fluorescence activated cell sorting (FACS, Figure 3C) using the Cas9-dsRed signal in order to enrich transfected cells (Figure 3D). Notably, no EGFP fluorescence signal could be observed after FACS. However, as sequencing results demonstrated successful integration, we concluded that the expression level of FE65 might be too low to be detectable by fluorescent microscopy.

**Figure 3:**
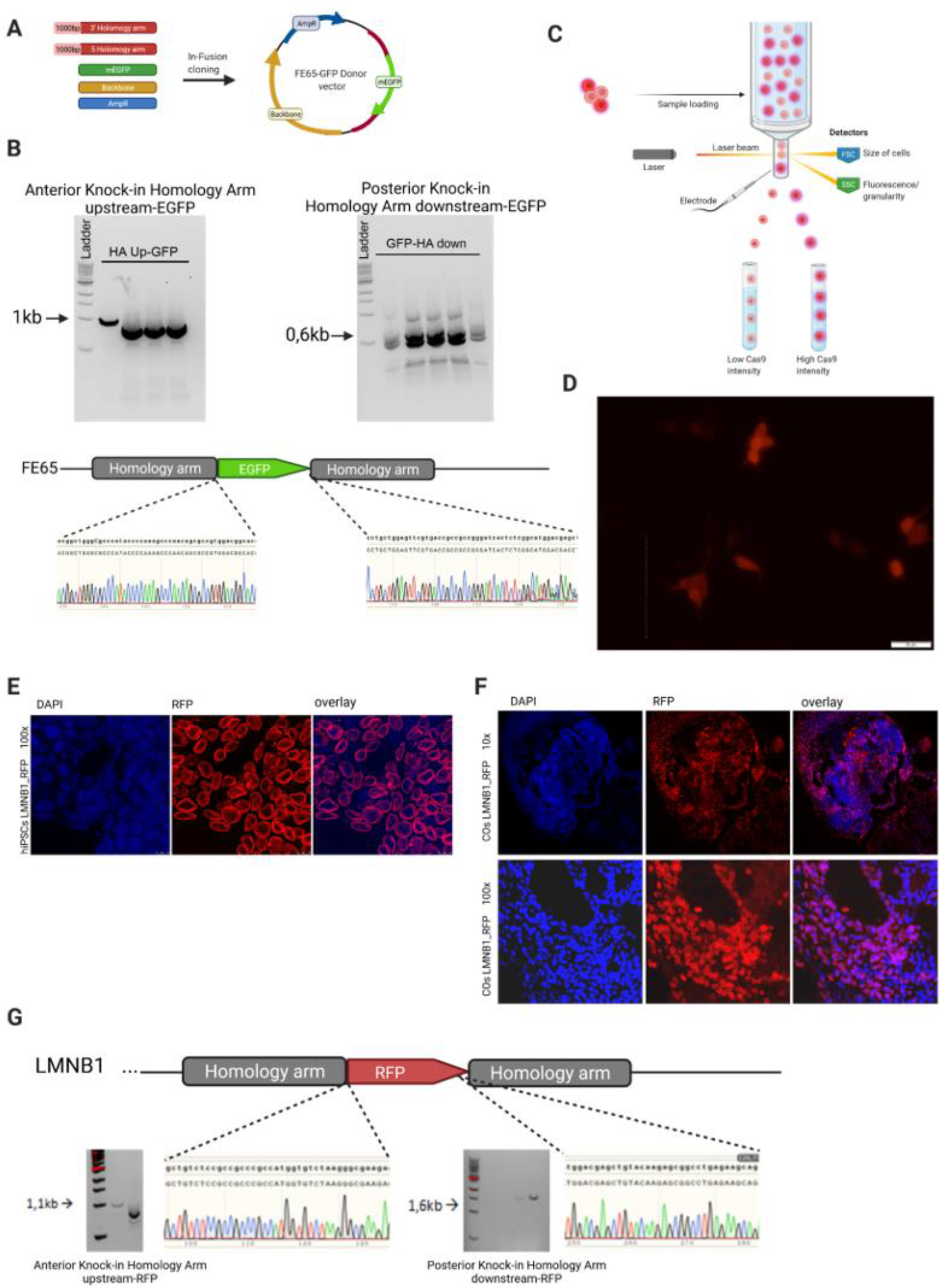
HDR-mediated approach demonstrated successful KI in HEK293T and hiPSCs. **(A)** The HDR vector was designed to contain the fluorescent KI reporter flanked by two homology arms of 1000 bp length (G-Blocks). **(B)** Upon transfection of Cas9, sgRNAs and the HDR vector, DNA isolation and PCR of the integration site, correct fragments for both, the anterior and posterior KI site were identified. Sequencing demonstrated seamless integration at both sites without any InDels. **(C)** Fluorescence assisted cell sorting (FACS) based on the Cas9-dsRED fluorescence was used afterwards. **(D)** Upon FACS, a 100 % dsRED-positive culture was obtained. No GFP signal (fused to the endogenous FE65) could be observed, pointing to low activity of the FE65 promoter not allowing fluorescence microscopy validation. (These results also confirm the assumption of false positive integration of the fluorescent cassette in the PCRext and NHEJ approach). **(E)** In order to validate the HDR-mediated approach and show robustness in other cell lines, we shifted to the highly abundant and subcellular characteristically localized LMNB1 target gene and used hiPSCs as another host cell line. HDR-mediated KI revealed the expected perinuclear RFP-fluorescent LMNB1 phenotype surrounding DAPI counterstained nuclei. **(F)** 10x and 100x magnification fluorescence microscopy pictures from 16 μm cryoslices of 21-days old RFP-LMNB1 cerebral organoids demonstrate the perinuclear ring-like LMNB1 phenotype in the 3D tissue. **(G)** Schematic representation of the N-terminal lamin B1 sequence, the location of the homology arms (1000 bp) and the RFP insertion site. Upon genomic DNA isolation, PCR amplification of the integration sites revealed distinct bands around 1.1 kbp and 1.6 kbp, respectively. Sequencing confirmed correct integration without InDels. (Figure created with https://BioRender.com)

### HDR-mediated KI in hiPSC and differentiation of gene-edited fluorescent organoids

Based on the successful results obtained in HEK293T cells, and in order to stress the effectiveness of HDR as a convenient KI method, we adapted the approach in order to tag the abundant protein lamin B1 (LMNB1) in hiPSCs. Therefore, we nucleofected hiPSCs with a donor plasmid containing LMNB1 homology arms flanking an RFP sequence. The cells that successfully included the KI were separated by FACS (Figure 3C) to receive a highly enriched culture. Due to the nuclear envelope localization of LMNB1, proper integration could first be verified by fluorescence microscopy as evident by characteristic fluorescent ring-like perinuclear structures (Figure 3E). PCR amplification from isolated DNA confirmed successful KI, as two fragments of the expected size (~1.1 kbp and ~1.6 kbp) were obtained. Sanger sequencing further proved the correct integration of the RFP gene in frame with the downstream and upstream homology arm sequence of LMNB1, notably without any InDels (Figure 3G). To exclude any impact of the editing approach the cellular pluripotency, RFP-LMNB1 hiPSCs were used to generate cerebral organoids. Fluorescence microscopy of cryosections of 21-days-old edited cerebral organoids revealed the characteristic LMNB1 phenotype, visualized as a red ring-like structure surrounding the nucleus in the differentiated tissue (Figure 3F). The HDR-mediated knock-in was the only approach demonstrating a completely correct KI that could be validated by extensive sequencing of the appropriate locus.

## Discussion

Our lab takes advantage of CRISPR/Cas9-mediated gene editing in order to knock-in large fragments for more than 5 years. Many strategies to do so have been suggested highlighting easy to use methods with fast and reliable results. On closer inspection, various publications rely on the pure fluorescence read-out, when knocking-in a fluorescence reporter, in order to confirm their work, without verification by DNA sequencing. Due to our own experiences and failings from carrying out some of the published methods, we aimed to compare the different approaches in order to identify the method of choice for the KI of large sequence fragments. Central requirement for this kind of experiment is the efficient cleavage of the genomic DNA. Indeed, all sgRNAs tested within this work demonstrated high cleavage efficiency, pointing to the robustness of the sgRNA/Cas9-mediated approach. The cloning-free generation of the donor template through simple PCR extension (PCRext) seems to be a promising, easy, and semi-fast method to generate the necessary donor fragment for the knock-in, which has been used in several studies (**Arbab & Sherwood, 2016; Arbab et al., 2015; Paix et al., 2015**). However, successful amplification strongly depends on primer quality (melting point, GC ratio, length), and in our case, it took several weeks to optimize PCR conditions, which contradicts the term “fast method” and rendered the method rather time-consuming. Further, there is the fact that in case of PCRext, there is low flexibility for the primer binding site driving amplification, which challenges the amplification optimization even more (different primer length, adaptation of the cycling conditions). Initially, we aimed to run three (instead of two) extension cycles in order to obtain a homology region >120 bp, however the third amplification failed, although several primer and cycle conditions were tested. Thus, we decided to proceed with the second extension corresponding to 120 bp homology length. Indeed, fluorescence analysis in transfected cells suggested a successful KI, but sequencing failed to validate correct integration. As the controls omitting the sgRNA were EGFP-negative (monitored for up to 7 days after transfection), these results point to an unspecific integration of the EGFP donor fragment, initiated by unspecific, sgRNA-driven cleavage in the HEK293T genome. The PCRext attempt indicates that a successful KI always needs to be validated by sequencing, and that relying on fluorescence signals alone is unreliable.

Another often used approach over the last 5 years is the NHEJ-mediated KI. NHEJ is a naturally occuring repair mechanism of the cell, known to be more efficient than HDR, but also being more error-prone (**Bétermier et al., 2014; Pannunzio et al., 2018; Song & Stieger, 2017**). The NHEJ repair mechanism has been utilized for KI experiments within the homology-independent targeted integration (HITI) approach, a method taking advantage of a linear donor fragment generated within the cell by sgRNA-mediated cleavage of a donor vector. Mostly, the donor vector is designed in a way that one of the sgRNA cleavage sites will bind to the endogenous integration site in parallel. In our hands, this approach was inefficient compared to the use of independent sgRNAs cleaving either the donor or the integration site, suggesting competing binding sites and that the exogenous donor vector might prominently recruit the sgRNA. Different sgRNA/donor vector ratios might overcome this limitation. While the fluorescence analysis of the cells transfected with our NHEJ-mediated design demonstrated a good efficiency of mNectarine expression, and therefore, indicated an effective knock-in of the donor fragment, the sequencing results pointed to a large variety of different InDels, which is characteristic for the NHEJ repair pathway. Thus, subsequent selection via FACS and cultivation of single cells is necessary to obtain monogenetic clones with InDel-free KI, which corresponds to a time-consuming and expensive workflow. Notably, for the posterior KI site, sequencing failed to show any correct integration suggesting that the number of cells that were correctly knocked-in at both ends of the strand tends to be very low. As for PCRext, the fluorescence analysis alone is not sufficient to verify the knock-in and NHEJ-mediated KI is not advisable for large fragment integration. Disillusioned by the results obtained so far, we next tested the HDR-mediated knock-in. Using standard cloning techniques, the vector design is challenging and time-consuming, as ~1000 bp homology arms for the sequence of interest have to be cloned and flanked to the KI fragment. Purchasing expensive G-blocks instead, massively facilitates this task, making HDR an interesting alternative approach for the KI of large fragments. In our hands, HDR-mediated KI was the only approach not causing non-specific fluorescence protein sequence integration as we observed for PCRext- and NHEJ-mediated KI. In other words, the missing fluorescence signal of the EGFP reporter, which could be successfully fused to the endogenous FE65 by HDR, reinforced our assumption of the unspecific integration within the PCRext- and NHEJ-based approach (where prominent EGFP signals were obtained, but sequencing failed to support correct integration). We assumed that the FE65 promoter activity might be too low to allow fluorescence imaging of the tagged protein, and thus decided to extend the study to the highly abundant and specifically perinuclear localized protein LMNB1. As demonstrated, this extension supported the HDR-mediated KI, which worked highly reliable in HEK293T cells as well as in hiPSCs and revealed no artificial InDels. Moreover, it was the only approach that could be reliably validated by comprehensive sequencing of the KI locus, re-emphasizing the necessity to not rely on pure fluorescence protein sequence integration. As true for all methods, the CRISPR/Cas9-driven approach might yield off-target effects, which was not in the scope of this work. As the approach could be extended to hiPSCs, the gene editing of the LMNB1 gene was confirmed by correct perinuclear localization and Sanger sequencing, and as edited LMNB1-RFP hiPSCs demonstrated full pluripotency to differentiate into cerebral organoids, we recommend HDR-mediated KI as a powerful and reliable tool to investigate protein function, localization, and protein-protein interaction *in live* in complex stem cell-derived 3D tissue.

**Table 1:** Overview of results from comparison of PCRext-, NHEJ- and HDR-mediated knock-in methods.

| Method | Advantages | Disadvantages |
| --- | --- | --- |
| PCR extension mediated | • less to moderate time-consuming, easy design<br>• cheapest method<br>• no cloning<br>• only short homology arms are needed | • ambitious PCR giving multiple side products<br>• time consumption rises with PCR efforts (multiple extension failed)<br>• low efficiency<br>• false positive results due to non-specific fluorescent protein integration<br>• sequencing failed to show KI<br>• not suitable for large KIs |
| NHEJ mediated | • allows knock-in in proliferating and non-proliferating cells | • time-consuming design<br>• highly error-prone, causing indels at insertion site<br>• intense downstream work: |
|  | • active in all stages of the cell cycle in a large variety of adult cells<br>• can be used in vitro and in vivo<br>• high efficiency because of direct linkage of DSB<br>• does not need homology arms, less time-consuming and labor-intensive, thus moderately cost-effective | culture and characterization of individual clones<br>• not suitable for large KIs |
| HDR mediated | • possibility of insertion of gain/loss of function gene (modeling of diseases)<br>• possibility of insertion of fluorescent tags in protein sequences,<br>• less error-prone than NHEJ<br>• only method working in our hands without InDels<br>• 100% correct sequencing results<br>• suitable for large KIs | • time consuming design and cloning if not G-Blocks are used<br>• expensive in case of G-Blocks used for vector design |

## Acknowledgements

This work was supported by funding from Deutsche Forschungsgemeinschaft (DFG) (MU3525/3-2), Mercur (Pr-2016-0010) and the Federal Ministry of Education and Research Germany (Bundesministerium für Bildung und Forschung; BMBF) (OrganSARS, 01KI2058). The instrument Leica TCS SP8 was supported by an instrument grant from the German Research Foundation (INST 213/886-1 FUGG).

---

**Supplementary Figure 1:**
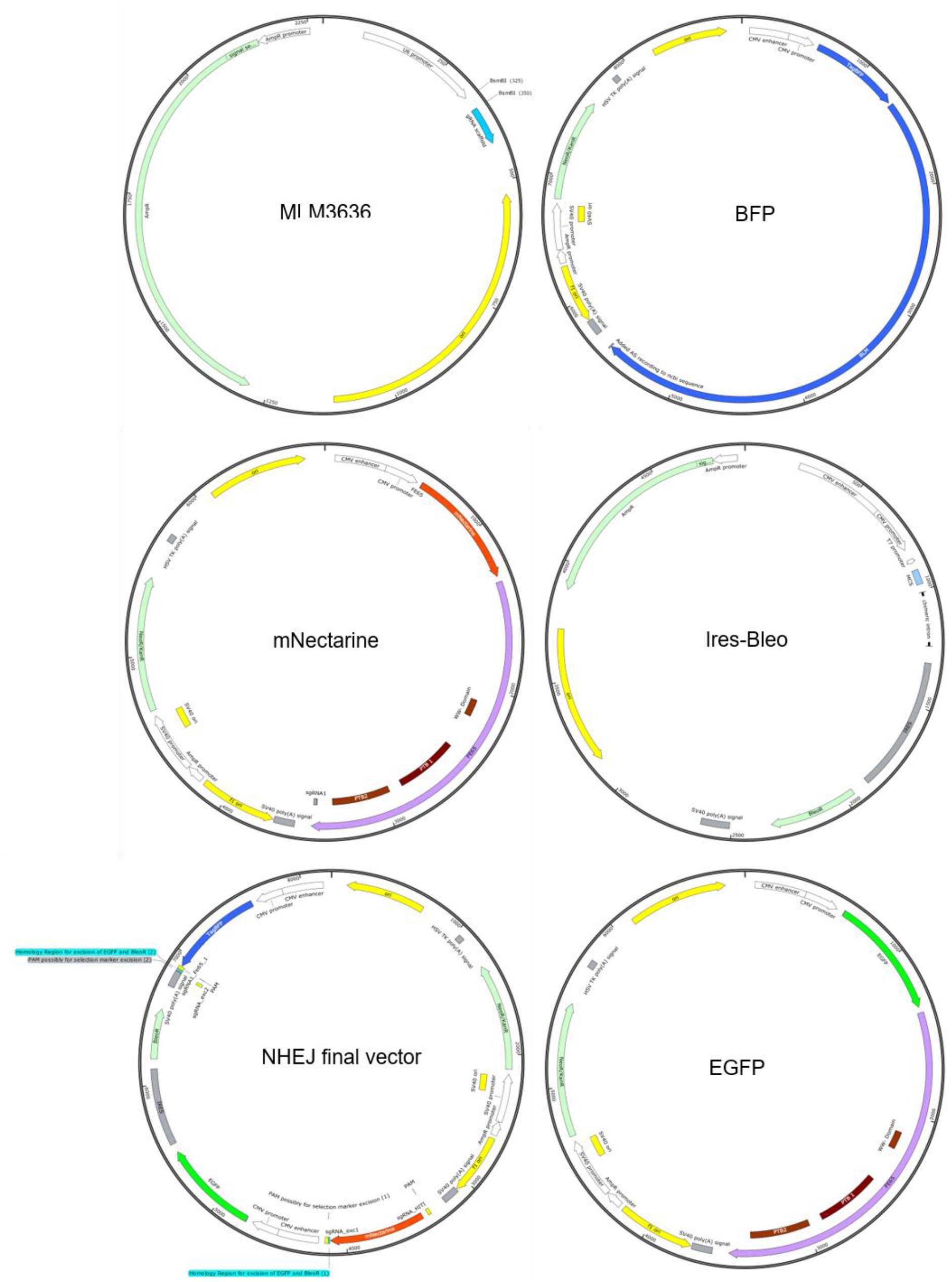
Overview of vector maps. Illustrated are vectors used for the generation of constructs in this paper. Maps were designed and visualized with Snapgene■. For sgRNA expression, the MLM3636 plasmid was used, while BFP, mNectarine, Ires-Bleo and EGFP were used to design and create the NHEJ final vector and FE65 GFP-Donor (HDR). Furthermore, the EGFP plasmid was used to generate the PCRext.

**Supplementary Figure 2:**
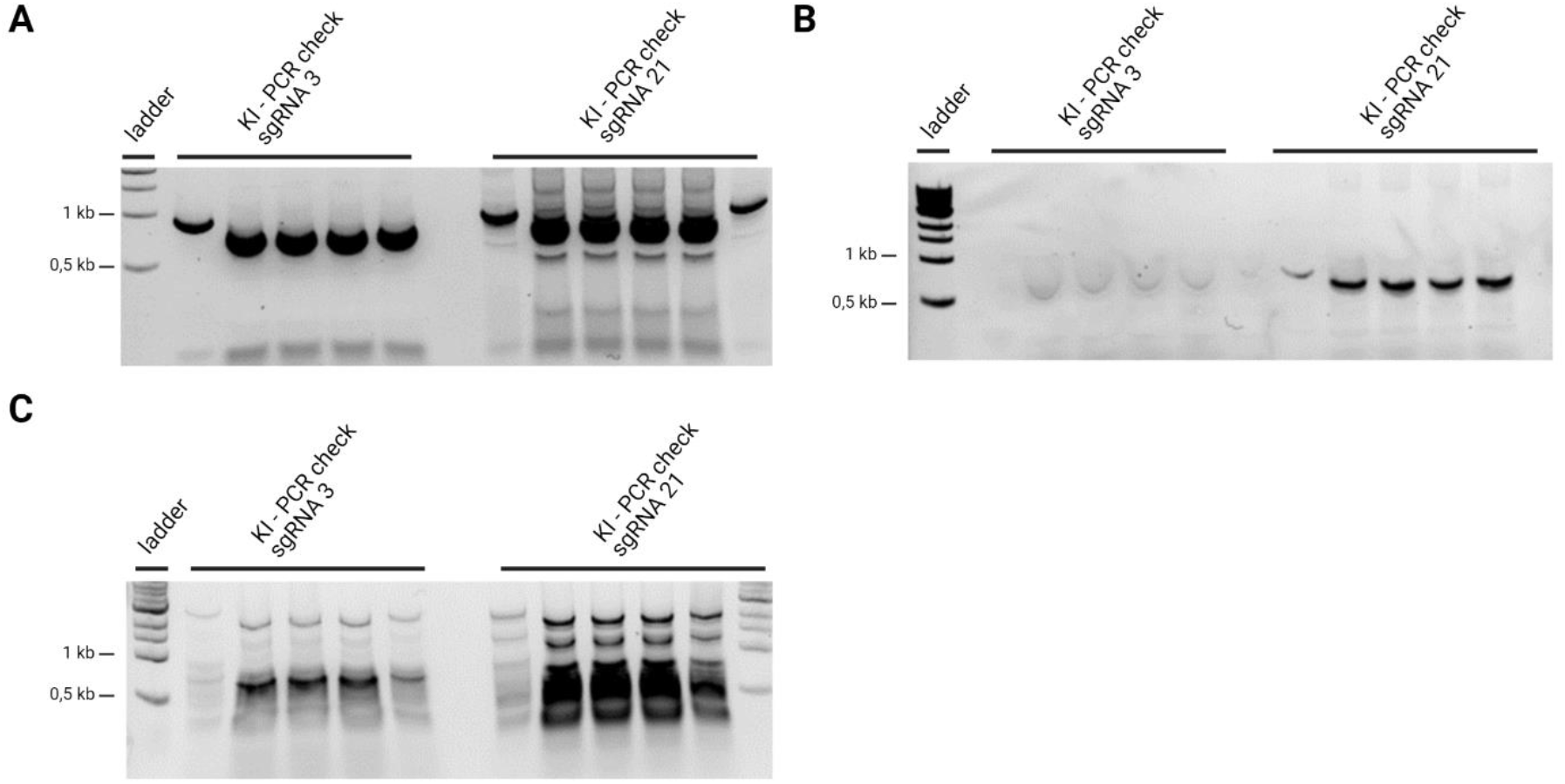
Gel electrophoresis results of PCRext study. **(A)** Primers were chosen to flank the knock-in site (HA upstream CATTGCACTGGGTTTATCCACTTACTG, HA downstream CAAAGACCCAAAGATACGTGGACAAAA). The expected band size with integrated EGFP was 1840 bp and would indicate a successful knock-in, while the band would be 1081 bp without the EGFP. Both bands were isolated. Sequencing failed. **(B)** First primer targeted in upstream HA (CATTGCACTGGGTTTATCCACTTACTG), while the second targeted the EGFP itself (EGFP rev TGGTAGAAGCCGTAGCCCATC). Expected product band size was 624 bp. The bands were extracted, sequencing failed. **(C)** EGFP for (GGTGGACAGCCACATGC) targeted within the EGFP and the second primer bond within the downstream homology arm (CAAAGACCCAAAGATACGTGGACAAAA) were used. The estimated product size was 831 bp, no sequencing results could be obtained.

